# Single-cell Multi-omics Integration for Unpaired Data by a Siamese Network with Graph-based Contrastive Loss

**DOI:** 10.1101/2022.06.07.495170

**Authors:** Chaozhong Liu, Linhua Wang, Zhandong Liu

**Affiliations:** Graduate Program in Quantitative and Computational Biosciences, Baylor College of Medicine, Houston, USA; Jan and Dan Duncan Neurological Research Institute at Texas Children’s Hospital, Houston, USA; Department of Pediatrics, Baylor College of Medicine, Houston, USA

## Abstract

Single-cell omics technology is being rapidly developed to measure the epigenome, genome, and transcriptome across a range of cell types. However, integrating omics data from different modalities is still challenging. Here, we propose a variation of the Siamese neural network framework called MinNet, which is trained to integrate multi-omics data on the single-cell resolution by utilizing graph-based contrastive loss. By training the model and testing it on several benchmark datasets, we showed its accuracy and generalizability in integrating scRNA-seq with scATAC-seq, and scRNA-seq with epitopes data. Further evaluation demonstrated our model’s unique capacity in removing the batch effect, which is a common problem in actual practice. To show how the integration impacts downstream analysis, we established model-based smoothing and cis-regulatory element inferring method and validated it with external pcHi-C evidence. Finally, the framework was applied to a COVID-19 dataset to compensate the original work with integration-based analysis, showing its necessity in single-cell multi-omics research.

## Background

Complex diseases like cancer, heart disease, and Alzheimer’s disease are caused by multiple unknown factors in the biological systems^1^. Many more units in various molecular layers play their roles in the disease progression than a simple Mendelian single-gene disorder, resulting in both aetiological and clinical heterogeneity. This heterogeneity complicates diagnosis, treatment, and the design and testing of new drugs^2^. With the development of high-throughput technologies, measuring multiple omics data such as scRNA-seq^3, 4^ and scATAC-seq^5, 6^ at the single-cell level has explained part of the heterogeneity from cell-type differences. However, a comprehensive and integrated view of all omics data is still lacking due to unpaired cells in different omics datasets. Thus, integrating different omics information is needed to elucidate potential causative changes that lead to disease, or the treatment targets, which can be tested in further molecular studies^7^.

Two main strategies have been proposed to integrate different omics data modalities: the experimental approach^8–11^, which profiles multiple omics data simultaneously on the same cells, and computational approaches, which fuses independent omics datasets. With the low throughput and high cost of experimental approaches^8^, the development of computational methods is of great importance to integrating multi-omics datasets. However, integrating omics data is still challenging due to the unpaired cells and modality/batch effect.

For unpaired cells problem, the general solution is to project all the cells into a shared nonlinear manifold or co-embedding space, from which unpaired cells can be aligned to share information of all omics data. Seurat^9^ applies canonical correlation analysis^10^ (CCA) to project datasets into common space and aligns cells by mutual nearest neighbors (MNN) for data fusion and label transfer. But it is criticized for using a linear dimension reduction algorithm and will distort the actual interrelationships between datasets^11^. Such linearity assumption was also adopted in Liger^12^, which uses integrated non-negative matrix factorization^13^. With the development of deep learning, an alternative to perform such nonlinear manifold projection is the encoder module from Autoencoder^14^. An encoder projects high-dimensional data into a low-dimensional representation with one or several layers of neurons. The succuss has been seen in GLUE^15^ using Variational Autoencoder.

The second challenge is technical effects from both modality and batch during integration.. Modality effect is removed when projecting and aligning cells in most algorithms. But to our knowledge, batch effect is not especially considered in these integration models. Previous work using the Siamese neural network^16^ has demonstrated the framework’s potential to integrate multiple scRNA-seq datasets and remove batch effects^17^. We believe such a Siamese neural network framework could also be constructed to integrate multi-omics data and removing both modality and batch effects. However, it is not applicable to use it directly to integrate datasets from multiple modalities at the single-cell level because it was trained to integrate single-modality RNA-sequencing data at a cell type level.

Therefore, here we introduce a new design of Siamese neural network with a graph-based loss to integrate multi-omics datasets at the single-cell resolution. Trained to integrate cells from different modalities while removing the potential batch effect, the model can complete the integration task while not biased by batch variation, showing superior performance in benchmarking datasets comparing other algorithms in multiple datasets. Furthermore, to show the impact of the integration on downstream analysis, we developed a model-based smoothing and cis-regulatory element inferring approach. We demonstrated its efficacy by validating in 10X Multiome datasets. Finally, the framework and analysis were applied to a published COVID-19 dataset, demonstrating its usability in compensating the integrated multi-modal analysis.

## Results

### Integrating single-cell multi-omics data through the Siamese neural network framework

The Siamese neural network receives simultaneously one cell from modality 1 (e.g., scRNA-seq) and another from modality 2 (e.g., scATAC-seq) as the inputs and projects them into a shared embedding space using the encoder. To make sure this manifold space is a good representation containing crucial biological information, two losses are applied following the encoder.

The first and the most important loss is the contrastive loss^18^. Contrastive loss aims to make similar cells close to each other and different cells separated in the joint embedding space. To achieve this goal, randomly chosen cell pairs are prepared before each training epoch for calculating either positive or negative contrastive loss. Positive pairs are corresponding cells in each modality (i.e., two modalities’ data of the same cell), and the loss is the Euclidian distance between the co-embedding spaces of the two modalities. Negative pairs are different cells sampled from the data, and the loss is calculated as a margin constant ***M*** minus the Euclidian distance. By training the model to minimize the loss, the distances between corresponded cells get smaller while the distances between negative pairs are larger.

Usually, the margin value ***M*** is a constant for all contrastive pairs during training. However, cells from different cell types are more diverse than cells from the same cell type. Thus, to make the integration work at single-cell resolution, we designed it as a flexible value depending on how different the cell pairs are in the graph we constructed before training (See **Method**). Intuitively, cells far from each other in the graph have a larger margin; highly similar cells close in the graph have smaller margin values in loss (**Figure 1B**).

**Fig. 1.**
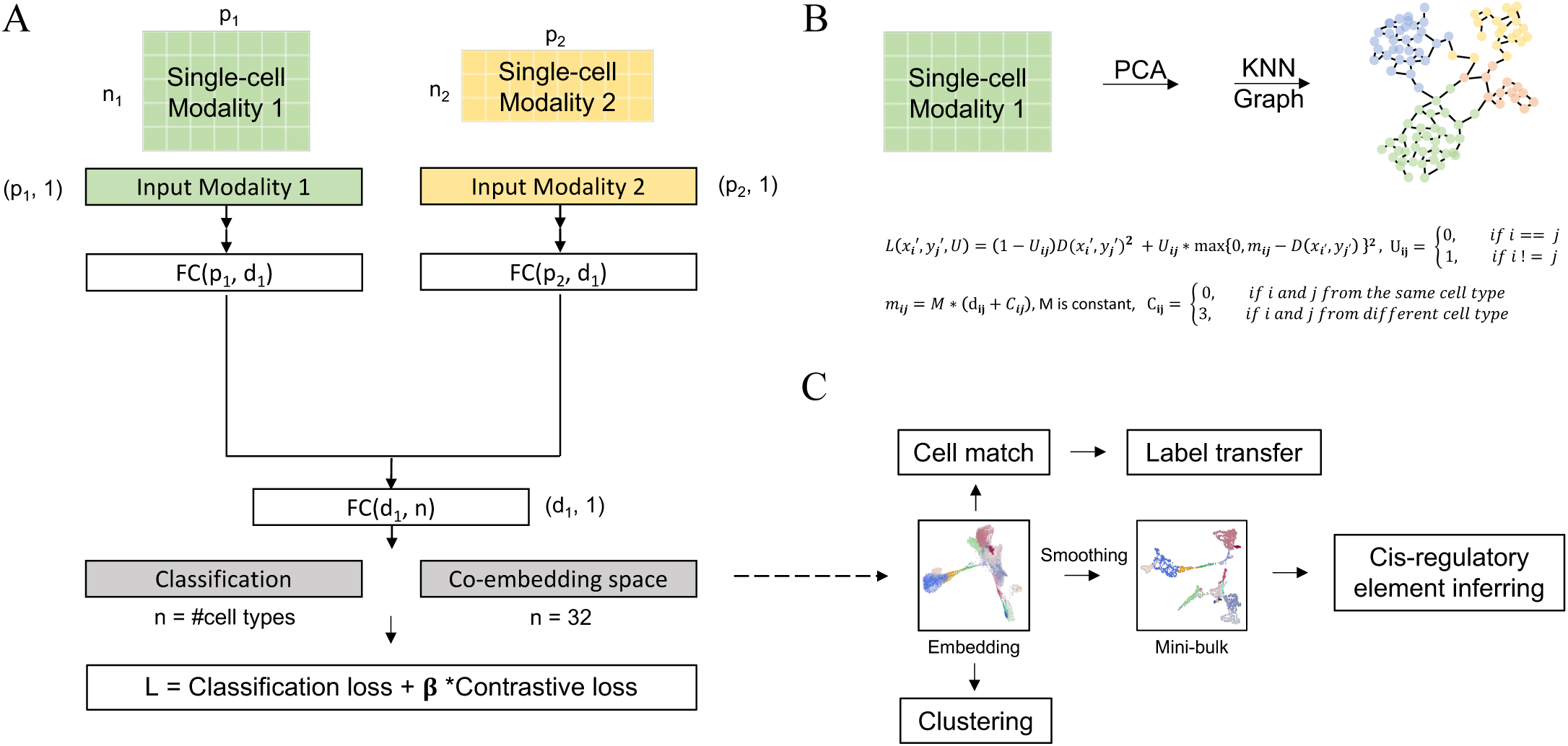
Overview of MinNet. **A** Model receives two modalities’ data as input. High-throughput omics data will go through an independent fully connected layer to be projected into a lower dimensional space. This representation space should be able to mix different modalities and separating cell types well. To achieve this, cell type classification loss and Siamese contrastive loss are used during training process. **B** To make the mixing resolution at single cell level rather than cell type level, we applied a KNN graph based Siamese loss with flexible margin value depending on cell pair graph distance. **C** In application, multiple omics data will be projected into this low-dimensional embedding space, based on which downstream analysis will be done, such as cell alignment, label transfer, unsupervised clustering, and the designed cis-regulatory element inferring pipeline.

**Fig. 2.**
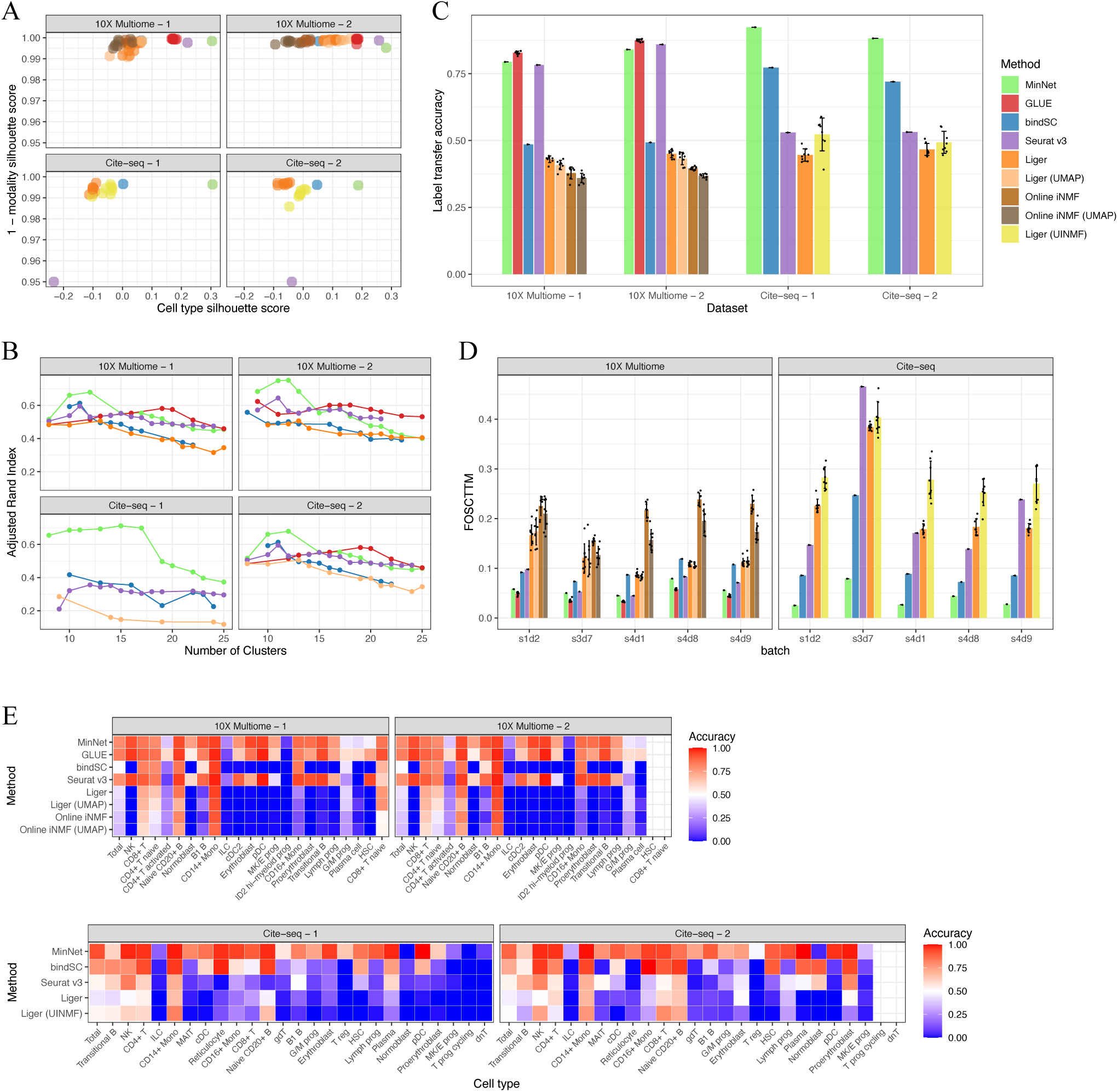
Performance benchmarks on gold-standard datasets. To test our model and compare with other existing algorithms, we benchmarked the transcriptome & chromatin accessibility data integration model and the transcriptome and cell-surface protein data integration model on datasets from the NeurIPS 2021 competition data. **A** Silhouette scores on the embedding space generated by all algorithms. Cell type silhouette score indicates how well cell types separate from each other, and 1– modality silhouette score indicates how well modalities are mixing with each other. **B** Adjusted Rand index along with the number of clusters comparing all algorithms. **C** Average label transfer accuracy bar plot. **D** FOSCTTM (Fraction of samples closer than the true match) score indicates the single-cell level alignment error of all algorithms. **E** Label transfer accuracy heatmap from transcriptome data annotation to either chromatin accessibility data or surface protein data.

The second loss is cell-type classification loss, which separates different cell types from each other. The output of the first encoder layer is sent to the label classification layer for cross-entropy loss calculation. An improved performance was previously observed in other studies thanks to its ability to accelerate the optimization process^17^.

The model needs to be trained with paired multi-omics datasets from techniques like 10X Multiome, SHARE-seq^19^, and SNARE-seq^20^, which profiles the transcriptome and chromatin accessibility simultaneously, or Cite-seq^21^, that profiles transcriptome and epitopes in the same cells. The weighted sum of classification and contrastive loss will be minimized during training to ensure optimized modality mixing and clustering.

We applied the framework in two tasks: transcriptome and chromatin accessibility data integration, which takes gene expression and gene activity score as input; transcriptome and epitopes data integration with gene expression and protein abundance. With the trained models, users can provide two simply normalized datasets and get the co-embedding space for downstream analysis, including aligning/pairing cells between modalities, unsupervised clustering, and cis-regulatory element inferring via pseudo-bulk generated from the embedding space.

### Benchmarking shows that MinNet is robust and generalizable in alignment and clustering

To test the performance and generalizability of our two models trained on 10X Multiome bone marrow mononuclear cells (BMMC) data and Cite-seq BMMC data, we evaluated our method and compared it with existing ones, including GLUE^15^, bindSC^22^, Seurat v3^9^, Liger^23^, and Liger’s online version^24^, on two untouched test sets from the NeurIPS 2021 Competition^25^ where the cell-to-cell correspondence is known. This large dataset has 10X Multiome and Cite-seq sequencing results from four sequencing sites and ten donors. Our models were trained on samples from part of the donors and three sequencing sites, leaving other donors untouched (test set 1) and the fourth sequencing site untouched (test set 2) as test sets. UnionCom^11^ failed all the evaluations due to memory overflow.

First, silhouette coefficient score^26^ was used to measure the integration performance of the co-embedding space generated by algorithms, regarding how well modalities are mixing while cell types are separating from each other. Compared with others, Siamese reached a higher score in both modality mixing and clustering (**Figure 1A**). The cell type silhouette coefficient indicates how well cell types are separated in the co-embedding space. In a real case, when researchers have no labels for their dataset, unsupervised clustering will be done to annotate the cell types. A better separation will ensure more precise annotations. We tested this by doing unsupervised clustering on the algorithms’ embeddings. The consistency between unsupervised clusters and cell type annotations was tested using the adjusted rand index^27^ (**Figure 1B**). Results show that MinNet-based clustering is the most concordant with the ground truth at the primary cell type level. And when subtypes were identified, our model was still competitive with the top ones in 10X Multiome data and is the best one in Cite-seq data. This cell type integration performance was also evaluated by the label transferring accuracy, which is another common task required in actual practice where researchers want to transfer the annotated labels from one modality to the other. Siamese can accurately transfer most of the labels, even if the cell numbers are small, while methods like Seurat are biased to the major cell types (**Figure 1C, E**). With Silhouette score and label transfer accuracy, our model’s performance is validated at the cell type level.

Beyond the cell type resolution integration, single-cell level cell alignment is also essential in some cases, like cell type sub-typing and mini-bulks generation for downstream analysis. To evaluate this higher resolution performance, FOSCTTM (Fraction of samples closer than the true match) score^28^ was measured for all co-embeddings generated. Siamese achieved the top 1 or 2 in all four datasets, especially in Cite-seq data, where it performed much better than the others (**Figure 1D**). These results showed our competitive performance at the single-cell resolution.

As a supervised model, the generalizability of it is essential to have a broader usability. The model’s generalizability has already been proven by showing the success of integrating untouched donor datasets and untouched sequencing site datasets. But we wanted to move on and test how this supervised model can be generalized to other tissues. First, the same evaluation was done on the 10X Multiome peripheral blood mononuclear cell (PBMC) dataset^29^. From the silhouette scores, FOSCTTM scores, and label transfer accuracies (Supplementary Figure 5A, C, D), our model still performed competitively compared with others and generated descent co-embedding space (**Supplementary Figure 6**). Though many of the cell types in the PBMC dataset were never learned by the model, it still separated most cell types well, demonstrating the model’s generalizability. Such generalizability is achieved thanks to the contrastive loss design, which was believed to learn the common sense of similar tasks rather than a specific task^30^.

However, the generalizability is limited to similar tissues, such as BMMC and PBMC. Furthermore, we applied the trained model to the 10X Multiome human brain dataset^31^, an entirely different tissue. As a result, the co-embedding space shows little biological information and failed to cluster well (**Supplementary figure 5B**). Therefore, we concluded that the generalizability of our algorithm could be expanded to similar tissues but not workable on distinct ones.

But this supervised approach is easy to be trained on targeting tissues and has a higher specificity, distinguish itself from others. For example, the traditional machine learning models, including bindSC, Liger, and Seurat, have better label transfer accuracies on the PBMC dataset than on the BMMC dataset. We believe it is due to cell type balance. When the numbers of cells in each cell type are pretty even, these methods perform well. But when it comes to cases like the BMMC datasets where monocytes take most of the whole dataset, label transfer of minor cell types based on these two methods is inaccurate. On the opposite, our approach is not influenced much by this unevenly distributed cell type number because of the superior specificity.

### MinNet shows the best performance in removing batch effect while maintaining biological variance

To distinguish between batch variance and biological variance, our model was trained with multiple batches from different donors and sequencing sites. While the input is normalized data without batch correction, the contrastive loss is based on the graph after batch correction by ComBat implemented in Scanpy^32^. Benefiting from this design, the model is required to produce the joint embedding that eliminates batch effects while keeping the biological differences.

We generated three more testing scenarios representing problems in the real case practice with the available benchmark datasets. The first scenario tests batch effect removal and clustering when both scRNA-seq and scATAC-seq experiments are performed independently on identical batches. The second and third cases are scenarios to test the integration performance of scRNA-seq and scATAC-seq datasets profiled from different batches. And again, we compared our model with the ones mentioned above.

The silhouette score and label transfer accuracy were evaluated only on the cell type resolution since cells are different in the second and third cases. **Figure 3A** shows the co-embedding space of the first case. While mixing the omics data, Siamese successfully separated cell types and mixed the batches. And the performance is quantified and compared among algorithms in **Figure 3B** using cell type and batch silhouette score. BindSC is better in mixing batches, but MinNetdistinguished the cell type variance from batch variance to give a better cell type separation. In the last two tests, most algorithms generated a co-embedding space that mixed the batch well, meaning all the manifold alignments between two single-batch omics data work (**Supplementary Figure 7, 8**). However, when it comes to the mixed batches, some algorithms failed due to their inabilities to take care of the batch effect, ending up with small cell type silhouette scores (**Figure 3B**). As a result, Siamese outperformed the others in clustering and label transferring (**Figure 3C**).

**Fig. 3.**
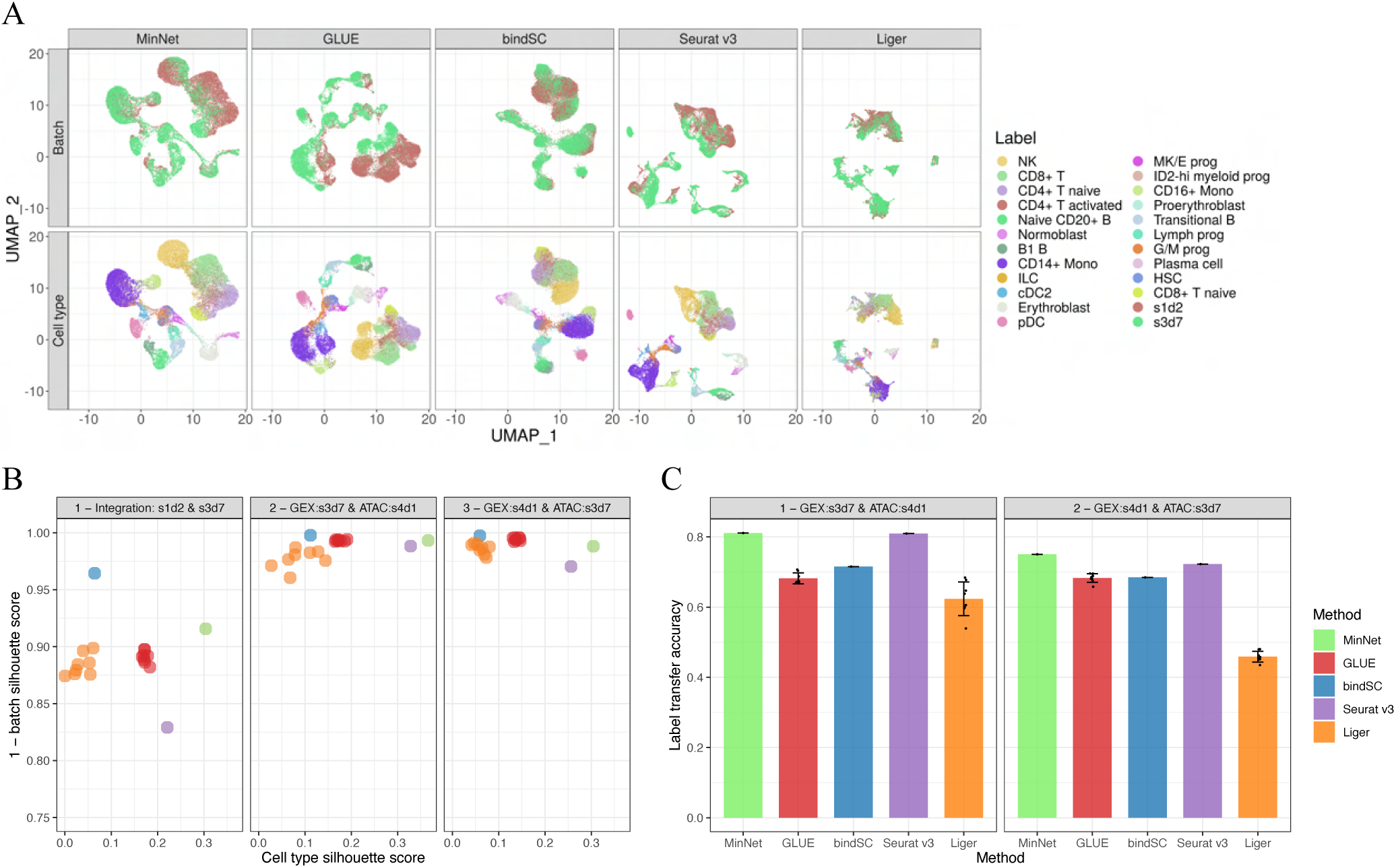
Batch effect removal of MinNet outperforms other algorithms. While separating cell types and mixing modalities, our model showed the best performance in removing batch effect, which is the most common problem met when integrating different omics data from distinct sources. **A** UMAP visualization of the embedding space generated by all algorithms. **B** Silhouette score indicates while separating cell types, our model mixes batches well. **C** Label transfer accuracy from one donor’s transcriptome data to another donor’s chromatin accessibility data.

We believe this evaluation is of more importance since it is close to the real practice where researchers got independent profiling from different batches or even from independent public available datasets. Testing on Cite-seq data with a similar case setting as the first one was also done (**Supplementary Figure 3**). We observed that none of the algorithms succeeded in mixing batches from different donors and sequencing sites, but they could mix batches from other donors and the same sequencing sites (**Supplementary Figure 4**). Thus, we hypothesize that it is due to the significant sequencing platform differences of Cite-seq technology. And more investigation should be done further.

### Model-based smoothing helps correlation-based cis-regulatory element inferring

After benchmarking our model, we demonstrate in the following writing part how integration can impact the downstream analysis and help discover biological findings.

To complement the high dropout rate and noise^33^ in single-cell data, we built functions to smooth^34^ (modifying matrix values according to their neighbor patterns) the data. To do this, we complemented the missing values in cells based on their K nearest neighbors in our single-cell resolution co-embedding space to decrease the sparsity (**Figure 4A**). After smoothing, mini-bulk data is generated before any downstream analysis such as cis-regulatory element inferring.

**Fig. 4.**
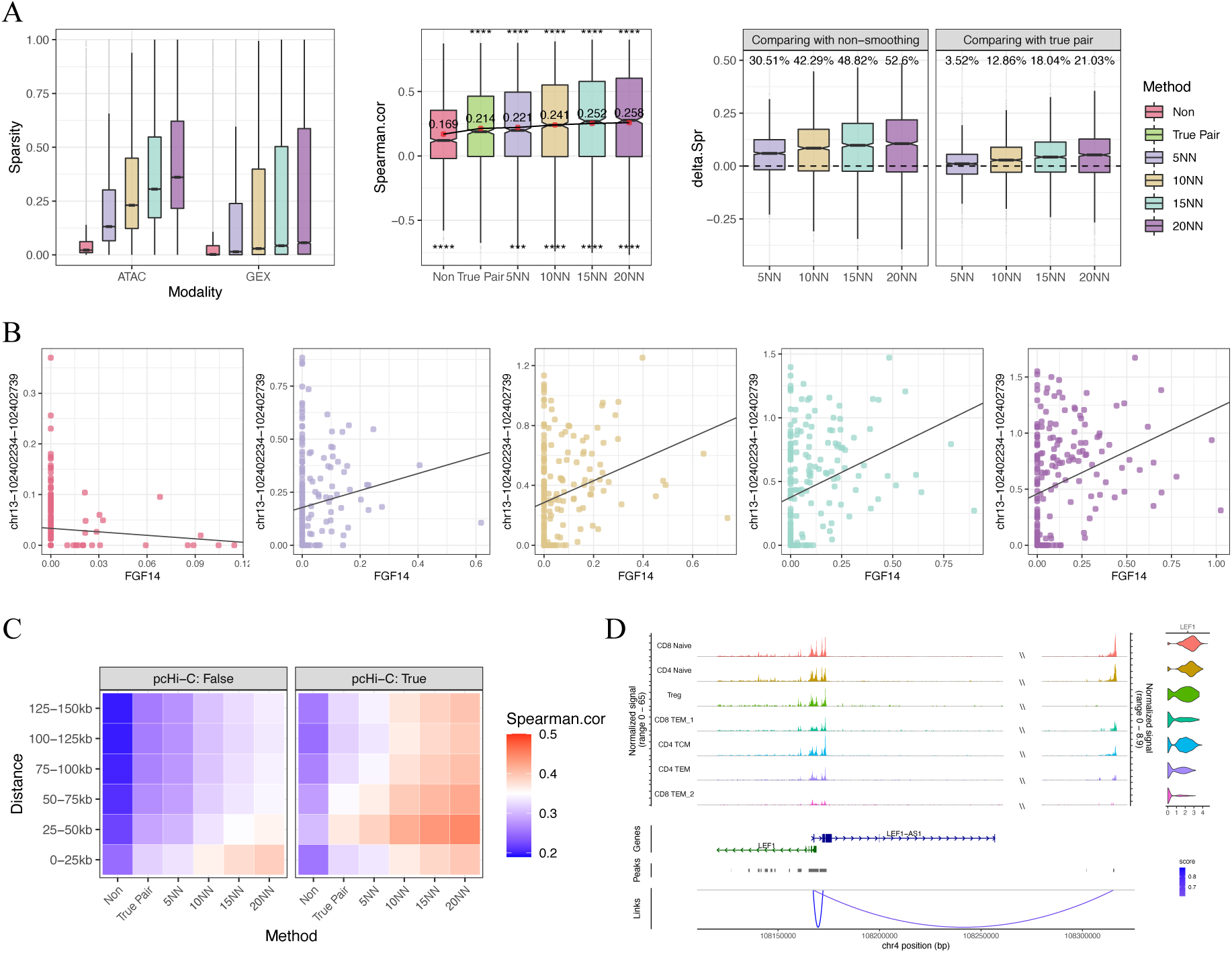
Model-based smoothing and cis-regulatory element inferring. By smoothing and generating mini-bulk omics profiles summing up neighborhood cells, we can infer the gene regulatory regions by calculating the correlation between transcriptome and chromatin openness. A higher correlation indicates a likely regulatory relationship between genes and peaks. **A** (Left) Smoothing decreased the sparsity of scRNA-seq and scATAC-seq data. (Middle) Smoothing increased the correlation between gene expression and its TSS regions openness compared with non-smoothing and true pair derived mini-bulk data. (Right) This trend is emphasized with showing the Spearman correlation coefficient differences between smoothed and non-smoothed mini-bulk data. **B** Example showing FGF14 and its TSS region peaks correlation in non-smoothing and smoothed data. **C** Heatmap showing the mean of correlation level between gene-peak pairs with different distances in all smoothed and non-smoothed datasets. **D** Genome track of LEF1 and its highly correlated peaks. Left shows the genome tracks of ATAC-seq data, right violin plots shows the gene expression level.

Cis-regulatory elements such as enhancers and promoters, are genomic regions that control development and physiology by regulating gene expression^35^. Inferring the regulation between open chromatin regions and gene expression is of great importance in understanding biological and disease process. Usually, cis-regulatory element inferring is done by calculating the correlation between chromatin regions, e.g., Cicero^36^. With integration done, we can now calculate the correlation between regions and gene expression using the paired cells or mini-bulks we aligned, which is more direct way in linking genome with transcriptome. Here, we implemented the smoothing, mini-bulk generating and cis-regulatory element inferring. The method was validated in the 10X Multiome peripheral blood mononuclear cells (PBMC) dataset^29^.

To test how this smoothing and mini-bulk benefit the downstream analysis, we calculated the correlations between genes and their 2kb nearby peaks in mini-bulk data, which are believed to be positively correlated. Results show that non-smoothing raw mini-bulk data has a lower correlation level than true pair mini-bulk data correlation, meaning that the dropout rate is a problem for downstream analysis when no cell correspondence is available between modalities (Figure 4A middle). But when smoothing was applied with the five nearest neighbors, the correlation level had already reached the ones of the true pair mini-bulk. The correlation was even higher than the true pair when increasing the number of neighbors. To demonstrate the importance of smoothing, we showed an example in Figure 4B. Chr3:102402234-102402739 is in the TSS region of the gene FGF14, which means the pair should be positively correlated. But because of the high dropout rate, non-smoothing mini-bulk data showed a negative Spearman correlation coefficient. When nearest neighbor complementation was applied, their association turned positive.

We further validated the model-based cis-regulatory element inferring approach with external pcHi-C evidences of the interaction between genome regions^37^. Non-smoothed, true pair, and smoothed mini-bulk data were applied to calculate the correlation between genes and their 150kb nearby peaks. While the mean correlations of pcHi-C unsupported peak-gene pairs weren’t increased much by smoothing, the mean correlation level of pcHi-C supported pairs was raised higher. And the difference in correlation between supported and unsupported pairs is clearly shown in the heatmap (**Figure 4C**). Again, with only five nearest neighbors smoothing, the correlation has reached the same level as true pair mini-bulk (Supplementary Figure 9A). But the 0-25k peak-gene pairs are non-distinguishable. We think it is due to the closeness to genes increased the co-openness even they don’t have a regulatory relationship.

Applying the model-based cis-regulatory element inferring, the extremely high correlation peak-gene pairs are worth further investigation because they indicate potential regulatory relationships. For example, LEF1 encodes the protein that can bind to a functionally important site in the T-cell receptor-alpha enhancer^38^. Thus, it shows a variant expression level in subtypes of T cells. Three peaks are within the 150kb upstream of the LEF1 TSS region. In 5NN smoothing mini-bulk correlation, 2 peaks have high correlations with LEF1 expression (chr4-108170508-108173850: r_s_ = 0.930; chr4-108315129-108315649: r_s_ = 0.877) and are supported by pcHi-C evidence. The other has a low correlation and is not supported by pcHi-C (chr4-108301923-108302013: r_s_ = 0.371). These results showed consistency with Hi-C and are validated by the data visualized in **Figure 4D**. On the other hand, some unsupported correlations also showed potential regulatory relationships. CCR2 encoded protein is a receptor for monocyte chemoattractant protein-1, a chemokine that specifically mediates monocyte chemotaxis^39, 40^. It is a monocyte marker; thus, the expression varies in many PBMC cell types. 6 peaks were shown high correlation with CCR2, 3 were supported by Hi-C evidence (chr3-46206526-46210451: r_s_ = 0.647; chr3-46297386-46301922: r_s_ = 0.612; chr3-46212074-46213996: r_s_ = 0.612) and 3 were not (chr3-46317953-46318717: r_s_ = 0.634; chr3-46312405-46313554: r_s_ = 0.607; chr3-46228191-46229079: r_s_ = 0.605). But when validated in the original data, we saw a correlation of all six peaks with the gene, indicating the potential genomic links (**Supplementary Figure 9B**). Furthermore, the unsupported chr3-46312405-46313554 together with Hi-C supported chr3-46206526-46210451 and chr3-46297386-46301922 were enriched in the motif for STAT3+IL-21 binding. IL-21 is a known cytokine with diverse effects on immune cells, including CD4 + and CD8 + T cells, B cells, macrophages, monocytes, and dendritic cells^41^. Thus, CCR2 might be involved in IL-21-induced cell adhesion through these binding sites, which was also discussed by other researchers^42^.

### MinNet compensates the lacking analysis of COVID-19 multi-modal data

To demonstrate how our model can help study diseases, we applied it to a publicly available COVID-19 dataset^43^ where healthy control and patients with various WHO severity score PBMC samples were profiled with independent scRNA-seq and scATAC-seq. We followed their scRNA-seq DEGs analysis and compensated for the missing part of the integration analysis to propose more potential discoveries. Two models were tried on the dataset: the PBMC trained model and the BMMC trained model.

Preprocessed and normalized data were provided to either PBMC or BMMC trained model to get the final co-embedding space (**Figure 5A**). Primary cell types are separating, and modalities are mixing well. Besides, even batch effect from different samples was removed, the severity difference is still clearly shown in the UMAP colored by the WHO severity score. Then we evaluated the consistency between the cell type annotation and our clustering by calculating label transfer accuracy from scATAC-seq to scRNA-seq or the reverse direction (**Figure 5B**). Except for cell types with only a few cells, the consistency is high between the two independent annotations and our embedding space. Both PBMC models and BMMC models were able to integrate the dataset due to their generalizability.

**Fig. 5.**
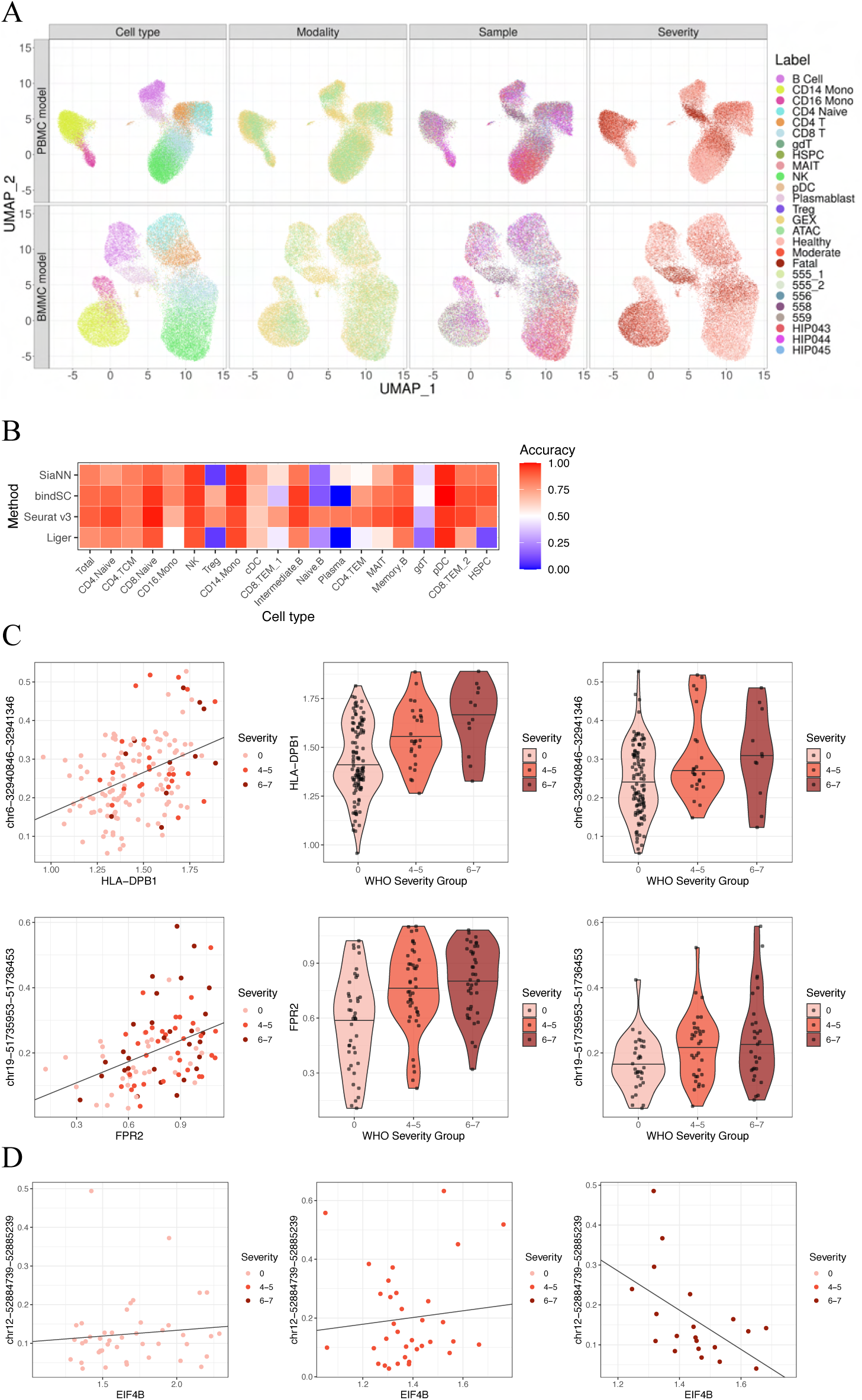
Application on Covid-19 dataset discovered cell type specific changes. Two models trained (BMMC and PBMC) were applied on this COVID-19 dataset. Health volunteers and patients PBMC transcriptome and chromatin accessibility were profiled independently to study the immune system changes due to the severity of Covid-19 infection. **A** UMAP visualization of the Covid-19 dataset labeled by cell type, modality, sample ID and severity. **B** Label transfer accuracy indicates that our embedding is consistent with the original cell type annotation on the majority cell types. Cell types with bad accuracy are due to only a few cells. **C** Example of regulatory element inferring from NK cells (upper) and Monocytes (lower). **D** Dysfunction of EIF4B might be due to the change in the regulatory role of correlated open regions.

With the co-embedding space output by the model, we generated the mini-bulk data per cell type. Then, we inferred the potential regulatory relationships between genes and peaks within a 150k bp distance from the transcription starting point. Two cell types, including NK cells and Monocytes, were chosen for the integration analysis because they are the mainly dysfunctional cell types concluded in the paper. Only DEGs by severity group were included in this analysis as compensation for their primary scRNA-seq DEG analysis. For example, in NK cells, the DEG HLA-DPB1 is correlated with chr6-32940846-32941346 (r_s_ = 0.333, Figure 5C) and showed differences among WHO severity groups. In monocytes, the DEG FPR2 is associated with the peak chr19-51735953-51736453 (r_s_ = 0.434, Figure 5C). These results were further validated with the raw data and could be potential regulatory sites for the DEGs.

Besides inferring the causal relationships between COVID-19 influenced peaks and genes, we can also compare the correlation among severity groups to discover dysfunction in severe groups caused by infection. Thus, we generated the pseudo-bulk data per cell type for each severity group separately and calculated the Spearman correlation between genes and peaks within a 150k bp distance. The inconsistency in correlation level might indicate the dysfunction between genes and peaks regulation. For example, in monocytes, EIF4B and its remote peaks chr12-52884739-52885239 are positively correlated in health control and moderate patients but are negatively correlated in severe patients (**Figure 5D**). Such findings indicate that the peak’s positive regulation is destroyed in severely infected patients. The potential rationale could be that the enhancing protein was competitively replaced by another inhibitory protein during the enhancer and promoter interaction, leading to the negative correlation between openness and expression.

## Discussion

We constructed this single-cell resolution multi-omics data integration model by designing the flexible margin contrastive loss based on the graph’s shortest distance. We successfully applied it to human BMMC scRNA-seq, scATAC-seq, and epitopes data integration. By benchmarking the performance, our model reached among the top methods comparing existing algorithms in silhouette score, FOSCTTM score, and label transfer accuracy. Benefiting from being trained in multiple batches and the loss design, the model can distinguish batch variation from actual biological variation and generate a better co-embedding space while mixing batches well. With the single-cell resolution and batch effect-removed embedding, better pseudo-bulk data can be generated for correlation-based cis-regulatory element inferring in integrating scRNA-seq and scATAC-seq. This whole model has been demonstrated in a real COVID-19 dataset to show how it can fill the gap in current multi-omics data analysis.

However, the drawback of our model is obvious too. This supervised model needs training on a multi-omics dataset from consistent or similar tissues with the needed ones to achieve the performance. Although we demonstrated the model’s generalizability by showing that our BMMC trained model can be successfully applied to different donors, different sequencing sites, and even PBMC tissues, the application to entirely different tissues still requires additional training on the specific tissue. However, we argue that such additional training is easily achievable. First, the number of paired single-cell multi-omics data is growing, providing sufficient tissue- and organism-specific training samples. Second, only a few hyper-parameters, including margin altitude ***M***, learning rate ***r***, and weight of contrastive loss, need to be tuned. Lastly, the training process is standardized and easily executable. Thus, applying the algorithm to a different dataset is an easy process. Nevertheless, we are working on more generalizable and unsupervised multi-modal integration models.

Another future improvement involves the gene activity score. It is known that this transformation of peaks will lose information^44^. And algorithms like GLUE and bindSC implemented their approaches to perform integration while optimizing the feature transformation between peaks and genes. Such design to maintain the peaks information will make the model more accurate, which will also be considered to improve our current MinNet model.

## Methods

### The Siamese neural network

The model receives simultaneously one cell from the single-cell modality one and another from the single-cell modality two as the inputs, denoted as 𝑥 ∈ ℝ^1×𝑝1^, 𝑦 ∈ ℝ^1×𝑝2^. P_1_ is the number of features in modality one, and P_2_ is the number of features in 2. The *x* and *y* will go through the encoding module first to get 𝑥′, 𝑦′ ∈ ℝ^1×𝐻^, H is the number of units in the hidden layer. Then, 𝑥′, 𝑦′ ∈ ℝ^1×𝐻^ will be linearly transformed into ℝ^1×𝑀^ vectors representing the probability of it belonging to each of the M cell types. Cross entropy loss is used for the final classification loss 𝐿_𝑙_ . Meanwhile, 𝑥′, 𝑦′ ∈ ℝ^1×𝐻^ will be linearly transformed to ℝ^1×32^ vectors representing its position on the final 32-dim joint embedding space. The contrastive loss will be calculated as follows:

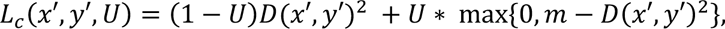

U is the label indicating whether the two cells are corresponding pairs (U=0) or not (U=1). D(.) defines the distance between 𝑥^%^ and 𝑦^%^. *m* here is the margin predefined between each pair of different cells using the shortest distance between the two in a KNN graph generated in preprocessing step (explained next). Intuitively, cells far away from each other in the graph has larger margin; highly similar cells that are close in the graph have smaller margin values (Figure 1B). Thus the total loss is:

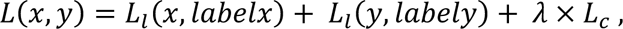

The *λ* is the weight between classification loss and contrastive loss.

### Determining the flexible margin from KNN graph

During training data processing, one of the omics data will be processed with batch correction, principal component decomposition (PCA) and KNN graph construction. The modality chosen is scRNA-seq for 10X Multiome and Cite-seq training data, because it presents the variation in data better in most of the cases. With the graph, the shortest distance ***d_ij_*** between all cell pairs will be calculated as part of the margin ***m_ij_***. The contrastive loss margin ***m_ij_*** of cell ***i*** from modality 1 and cell ***j*** from modality 2 is defined as:

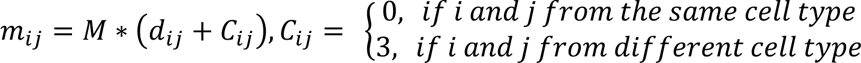

M is a constant controlling the scale of contrastive loss and can be the tunable hyper-parameter. C is used to increase the penalty of two cells of different cell types being close to each other in the co-embedding space, value 3.0 worked in all our scenarios.

### Training process

The two preprocessed feature matrices are scaled by genes to unit variance and zero mean, followed by clipping values larger than 10. We use the Adam optimizer to train the model with user-provided hyper-parameter values including M, *λ*, and learning rate. Before each epoch, the two matrices will be shuffled, and all cells are randomly assigned either a positive (same cell in different modalities) or negative (different cells) cell pair to calculate the contrastive loss. The number of negative pairs and positive pairs is controlled near 3:1.

Two assigning strategies are tried and perform equally well. First is the between-modality strategy, negative pairs are different cells in different modalities. Second is the within-modality strategy, negative pairs are different cells in the same modality. We believe it is due to positive pairs being the same for the two strategies so both within and between-modality co-embedding space correction works well. In the final model, we chose the between-modality strategy for scRNA-seq and scATAC-seq integration task, and the within-modality strategy for scRNA-seq and cell surface protein integration task.

### Data processing

The preprocessed and well-annotated bone marrow mononuclear cells data from the NeurIPS 2021 competition can be downloaded in **GSE194122**. The AnnData object was loaded in python 3.6.13 with AnnData 0.7.6. All Scanpy based processing mentioned below is done with Scanpy 1.7.2.

### NeurIPS 2021 competition 10X Multiome training data

Samples from sequencing sites 1 and 2 were taken as the training dataset including s1d1, s1s3, s2d1, s2d4, s2d5, s3d3, s3d6, and s3d10. S stands for sequencing site and d stands for donor number. First, we performed feature and cell selection. Highly variable genes in scRNA-seq data of all batches were determined by Scanpy *pp.highly_variable_genes* function with Cell Ranger flavor. Only genes marked as highly variable genes in more than 1 batch were kept. Besides, we also kept cell surface protein genes as the features for training. Then, we performed more strict cell filtering based on mitochondria gene expression proportion (<4), number of genes expressed (100 - 4000), number of peaks (1000 - 80,000), and the total number of fragments (1000 - 300,000). This is the final feature set and cell set for training.

We used the already processed gene activity matrix saved in the Anndata obsm *gene_activity*. It is the count sum of peaks 2kb upstream of the selected genes’ TSS region calculated by Seurat v3. Together with the feature-selected scRNA-seq data, log-transformed per cell normalization is performed to correct sequencing depth difference. These are the final input of two matrices for model training.

To determine the margin value between cell pairs, we constructed a KNN graph using the scRNA-seq data. ComBat implemented in Scanpy was used to perform batch correction, followed by PCA and K nearest neighbor graph construction saved as a large sparse matrix in the AnnData object named *connectivity*. Distances between neighbor cells were then estimated by 1.01 - *connectivity* value. To calculate the shortest distance between all pairs from the large sparse matrix efficiently, Scipy 1.5.4 *dijkstra* function was used to generate the NxN matrix recording all shortest distances for training.

### NeurIPS 2021 competition Cite-seq training data

Samples from sequencing sites 1 and 2 were taken as the training dataset including s1d1, s1d3, s2d1, s2d4, s2d5, s3d1, and s3d6. We performed similar feature and cell selection as 10X Multiome data in GEX data. Highly variable genes in scRNA-seq data of all batches were determined by Scanpy *pp.highly_variable_genes* function with Cell Ranger flavor. Only genes marked as highly variable genes in more than 2 batches were kept. Besides, we also kept cell surface protein genes as the features for training. Then, we performed more strict cell filtering based on mitochondria gene expression proportion (<15), the number of genes expressed (75 - 1200), and the number of peaks (75 - 1500). All features in ADT data were kept and cells are kept the same as GEX data. The final feature and cell-selected matrices were under log-transformed normalization before training.

The same strategy was applied as 10X Multiome dataset to determine the shortest distances between all cell pairs.

### NeurIPS 2021 competition 10X Multiome test data

Samples from s1d2 and s3d7 were taken as the first testing set. Samples from s4d1, s4d8, and s4d9 were taken as the second testing set. Log-transformed transcriptome matrix is used as one of the inputs for the trained model. The gene activity matrix is from the already processed ones saved in the Anndata obsm *gene_activity*. Before applying the neural network, we selected the features in training and compensate for missing features with all 0 values. Then the two matrices were scaled by genes to unit variance and zero mean, followed by clipping values larger than 10. Finally, the test mode of the trained model was run to generate the 32-d co-embedding space coordinate for every cell.

### NeurIPS 2021 competition Cite-seq test data

Samples from s1d2 and s3d7 were taken as the first testing set. Samples from s4d1, s4d8, and s4d9 were taken as the second testing set. The same process is done as 10X Multiome data besides we used the ADT data instead of the gene activity matrix.

### Human peripheral blood mononuclear cells (PBMCs) Multiome data from 10X Genomics

The dataset can be downloaded on 10X Genomics website at https://support.10xgenomics.com/single-cell-multiome-atac-gex/datasets/1.0.0/pbmc_granulocyte_sorted_10k. We followed all the same processing of Seurat integration tutorial document at https://satijalab.org/seurat/articles/atacseq_integration_vignette.html. The gene activity matrix was calculated using Signac 1.1.1 summing up counts 2kb upstream of the gene TSS region. Gene activity matrix and genes count matrix were saved as HDF5 files together with the metadata. Then the files were loaded in python and went under log-transformed normalization using Scanpy. The following processes were the same as NeurIPS 10X Multiome data.

### Human brain Multiome data from 10X Genomics

The dataset can be downloaded on 10X Genomics website at https://www.10xgenomics.com/resources/datasets/frozen-human-healthy-brain-tissue-3-k-1-standard-2-0-0. Preprocessing was done by filtering cells in RNA-seq that have less than 1000 counts, larger than 25000 counts or high mitochondria proportion (>10%), and filtering cells in ATAC-seq with fragment counts less than 5000 or larger than 1e5. Only cells remaining in both modalities were kept. Dimension reduction and clustering were done following Surat default pipeline. The gene activity matrix was calculated using Seurat v3 summing up counts 2kb upstream of the gene TSS region. Gene activity matrix and genes count matrix were saved as HDF5 files together with the metadata. Then the files were loaded in python and went under log-transformed normalization using Scanpy. The following processes were the same as NeurIPS 10X Multiome data.

### JEM COVID-19 Multi-omic profiling scRNA-seq data

The fully processed scRNA-seq AnnData H5AD file can be downloaded at https://www.covid19cellatlas.org/index.patient.html. Metadata is downloaded at their GitHub page at https://github.com/ajwilk/COVID_scMultiome. We performed log-transform normalization with the processed data. And to keep consistent with scATAC-seq data, we kept only shared donor batches and re-annotated cell types.

### JEM COVID-19 Multi-omic profiling scATAC-seq data

The raw data can be downloaded at **GSE174072**. The fragment files were processed using ArchR 1.0.1 following the same quality control as mentioned in the paper. The batch-specific TSS enrichment score and the minimum number of fragments cutoff can be found on their GitHub page mentioned above. Worth mentioning here is though they claim the sequencing reads were aligned with hg19 reference genome, we found only using hg38 can give the correct TSS enrichment score. Thus, hg38 was used in all following related processes. We followed the same data processing pipeline in the paper, removing doublets, clustering, batch correction with Harmony, and calling peaks with MACS2. To follow the same practice as the training dataset, we used the Seurat 3.1.1 *CreateGeneActivityMatrix* function to generate the gene activity matrix instead of using ArchR provided gene activity matrix. The saved HDF5 file was loaded in python and compiled into AnnData object together with the metadata from their GitHub page and went through log-transformed normalization. Again, to keep consistent with scRNA-seq data, the cell type was re-annotated, and only shared batches were kept. Finally, the two log-transformed matrices were provided to the model followed the same pipeline as other test datasets.

### Running of all algorithms

GLUE 0.1.1, bindSC 1.0.0, Seurat 3.1.1, UnionCom 0.2.3, Liger 1.0.0, its Online-iNMF and UINMF version were all included to get a systematic benchmarking. Due to the memory outflow problem, UnionCom failed to get the results with GLUE-provided codes on their GitHub page. All others were implemented successfully according to the authors’ tutorial. All codes are available at GitHub.

We followed GLUE’s tutorial at https://scglue.readthedocs.io/en/latest/tutorials.html with all default settings. We started from raw test data to run the data preprocessing and model training steps. The final cell co-embedding space is saved for all benchmarking. GLUE was run 8 times with different random seeds.

We followed bindSC’s tutorial at https://htmlpreview.github.io/?https://github.com/KChen-lab/bindSC/blob/master/vignettes/CITE-seq/CITE_seq.html for Cite-seq data integration and https://htmlpreview.github.io/? https://github.com/KChen-lab/bindSC/blob/master/vignettes/method_eval/method_eval.A549.html for 10X Multiome data integration task. The final embedding used was the bi-CCA generated results. BindSC was run 8 times with different random seeds.

We followed Seurat’s tutorial at https://satijalab.org/seurat/articles/atacseq_integration_vignette.html for both Cite-seq and 10X Multiome data integration. The final embedding space is the UMAP dimensional reduction space following Seurat integration pipeline.

Liger and its online iNMF version were implemented for 10X Multiome data integration, following the tutorials at https://htmlpreview.github.io/?https://github.com/welch-lab/liger/blob/master/vignettes/walkthrough_rna_atac.html and http://htmlpreview.github.io/?https://github.com/welch-lab/liger/blob/master/vignettes/online_iNMF_tutorial.html. Cite-seq data was integrated using Liger and its UINMF version at http://htmlpreview.github.io/?https://github.com/welch-lab/liger/blob/master/vignettes/UINMF_vignette.html. Each model was run 8 times with different random seeds.

All the UMAP visualizations were done either using the software available functions or using Scanpy default settings.

### Benchmark criteria

Silhouette score was used to evaluate how well cell types are clustered and modalities are mixed. It is a measure of how similar an object is to its own cluster (cohesion) compared to other clusters (separation). The silhouette ranges from −1 to +1, where a high value indicates that the object is well matched to its own cluster and poorly matched to neighboring clusters. The silhouette score was calculated using Scikit-learn 0.24.2. To measure how well cell types are clustered, the raw silhouette score value was used. And for modality mixing and batch mixing, 1 - silhouette value was used. So the higher the score, the better the performance.

Rand index is a measure of the similarity between two data clusterings. Adjusted Rand Index was calculated using Sklearn 1.0.1 *adjusted_rand_score* function comparing unsupervised clustering and cell type annotations. Unsupervised clustering was done using Scanpy *tl.leiden* function with different resolutions so that all algorithms can get the evaluation on cluster numbers from 8 to the number of cell types in each dataset.

FOSCTTM (Fraction of samples closer than the true match) score was used to evaluate the co-embedding space at single-cell resolution. Assume two single-cell omics data profiled the same set of N cells, when cells are projected into the co-embedding space, the FOSCTTM we calculated is defined as:

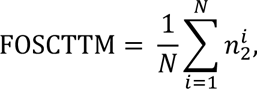

where 𝑛^+^ means the number of cells in the second modality that are closer to the i^th^ cell in the first modality than its true matches in modality 2.

Label transfer accuracy is used to measure the performance of all co-embeddings on this common task. The transfer is measured from scRNA-seq cell type annotations to either scATAC-seq or cell surface protein data. Except Seurat using their own label transfer method, all other algorithms’ label transfer is done by weighted K nearest neighbors. It means the label of cell from the second modality is predicted as the max weighted vote of its K nearest cells in scRNA-seq. K is chosen for each algorithm when it reached the best performance. With the predicted cell type label and the true label, the label transfer accuracy is defined as:

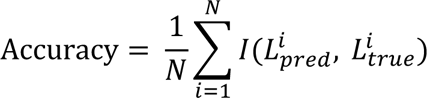

### Data Smoothing, mini-bulk generating and Cis-regulatory element inferring

Correlation-based regulatory element inferring is always weakened because of the high dropout rate. To solve this, we first do data smoothing and generat transcriptome and chromatin accessibility mini-bulk data with the single-cell resolution co-embedding space.

#### Smoothing

Nearest neighbor graph was constructed based on the model generated co-embedding space using Scanpy *pp.neighbors* function with default parameters and different number of neighbors including 5, 10, 15 and 20. The raw count matrix was multiplied by the binarized connectivity matrix to complement missing values by neighbors. The connectivity matrix was binarized by two steps: 1) cells are the nearest neighbors of the target cell; 2) the distance between them should be smaller than the 95% distance percentile value. Cells passing the criteria will be used to complement the target missing values by keeping the value as 1 in the binary connectivity matrix.

#### Mini-bulk

We used Scipy 1.5.3 *pdist* function to perform hierarchical clustering of all cells in the two modalities. Then with the hierarchical order and cell types, cells were cut into N = 100 mini-bulks, and each mini-bulk is ensured to contain only one cell type. The mini-bulk matrix is generated with scglue 0.1.1 *aggregate_obs* function.

Next, genes of interest were selected and the correlation of them with their 150kb upstream peaks were calculated from the mini-bulk data. The abnormally high correlations between remote peaks and genes might indicate the cis-regulatory relationships. Correlation results were visualized with box and violin plots using ggplot2 in R.

### pcHi-C data processing

pcHi-C data is available at https://ars.els-cdn.com/content/image/1-s2.0-S0092867416313228-mmc4.zip and https://osf.io/e594p/. Our codes were based on GLUE’s processing and scglue functions but simplified to only extract the pcHi-C evident pairs. Only evidence of overlapped cell type were chosen to be validated. Then scglue is used to map these peak-gene pairs to the 10X Multiome PBMC dataset peak-gene pairs. The evidence is save as .graphml file for read and write efficiency.

### Enrichment analysis with Homer

Peaks of interest were listed in BED format as the input for Homer v4.11.1. Function *findMotifsGenome* were applied to enrich the peaks of interest in known motifs. For some of the cases, we have only a few peaks, so the statistical test is not reliable. In cases like that, we only trust the information about which motifs are matched with most of the peaks.

### Data Visualization

All visualization figures were done using ggplot2. The example genome tracks were plotted with ArchR *plotBrowserTrack* and Seurat v3 *CoveragePlot* function.

### Code Availability

All codes are available on **GitHub** at https://github.com/ChaozhongLiu/MinNet.

## Notes

### Competing Interest Statement

The authors have declared no competing interest.

### Summary of Updates

We changed some typo about the framework name.

