## Supplementary figures and images for "Single-cell Multi-omics Integration for Unpaired Data by a Siamese Network with Graph-based Contrastive Loss" (from another version of this preprint: version 1, 2022-06-08)

### Supplementary Figure 1

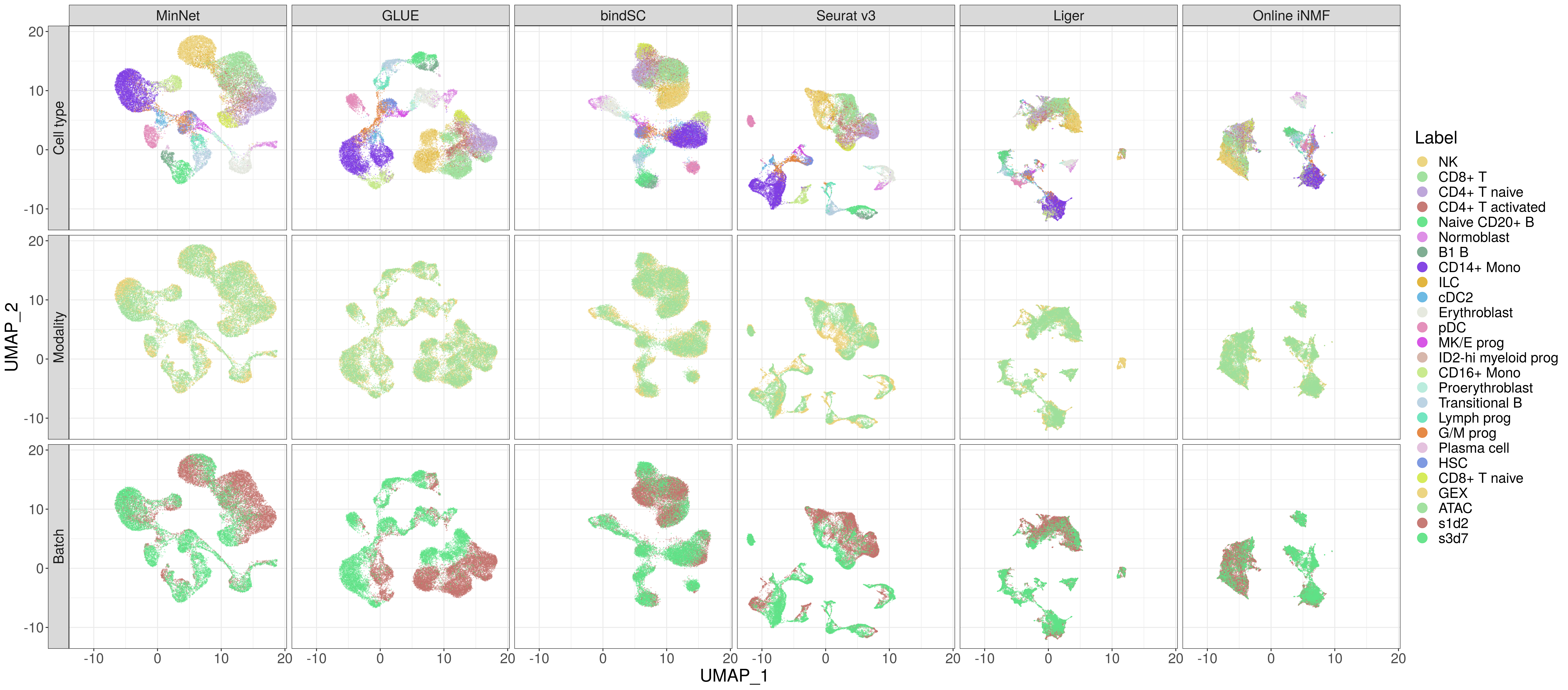

### Supplementary Figure 2

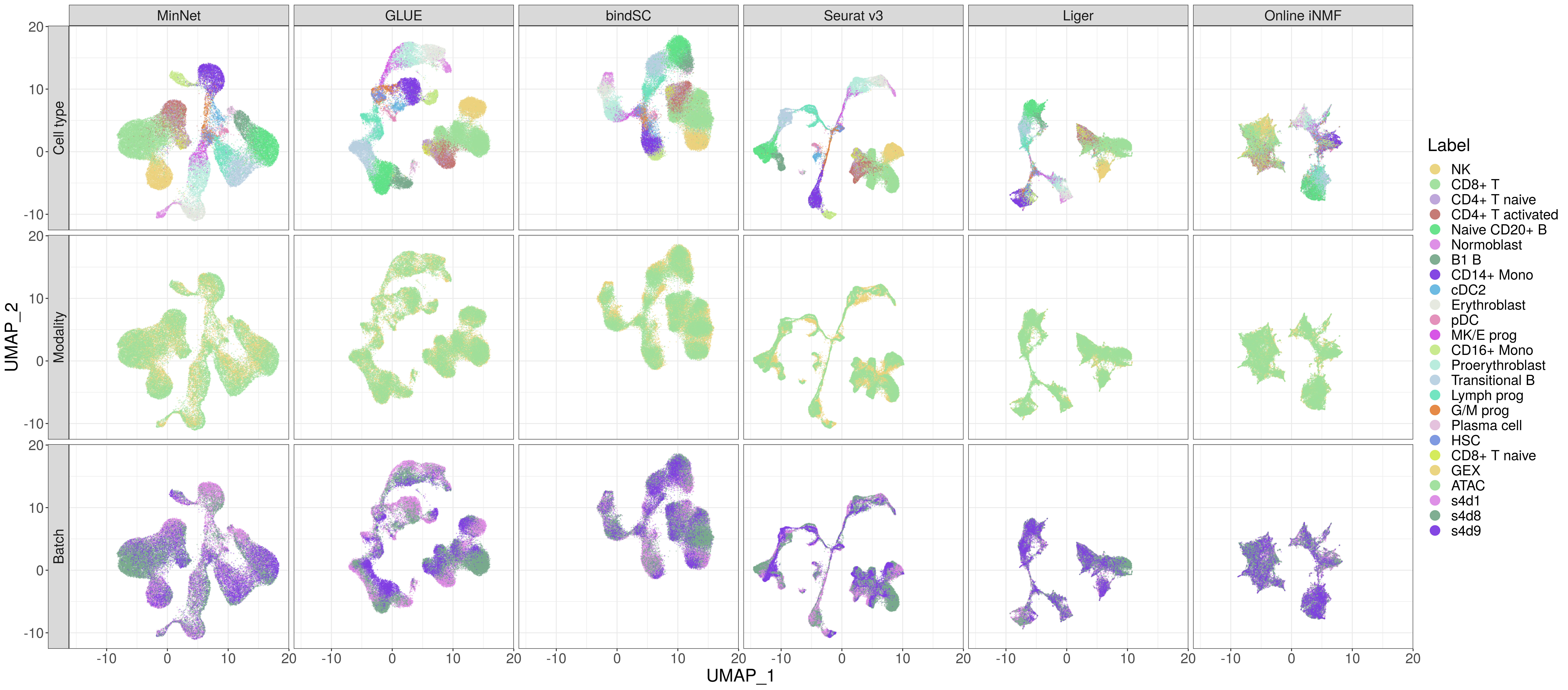

### Supplementary Figure 3

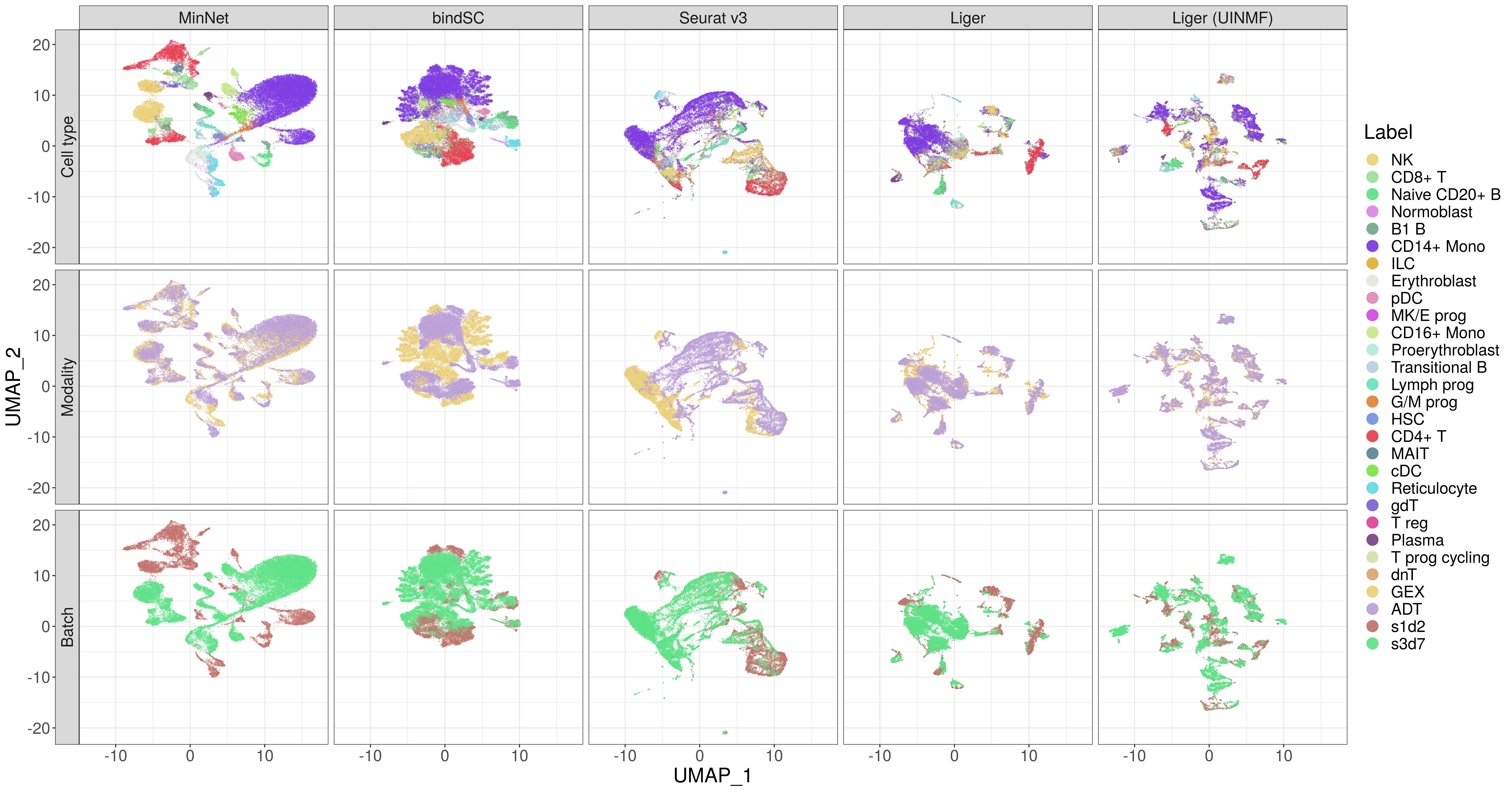

### Supplementary Figure 4

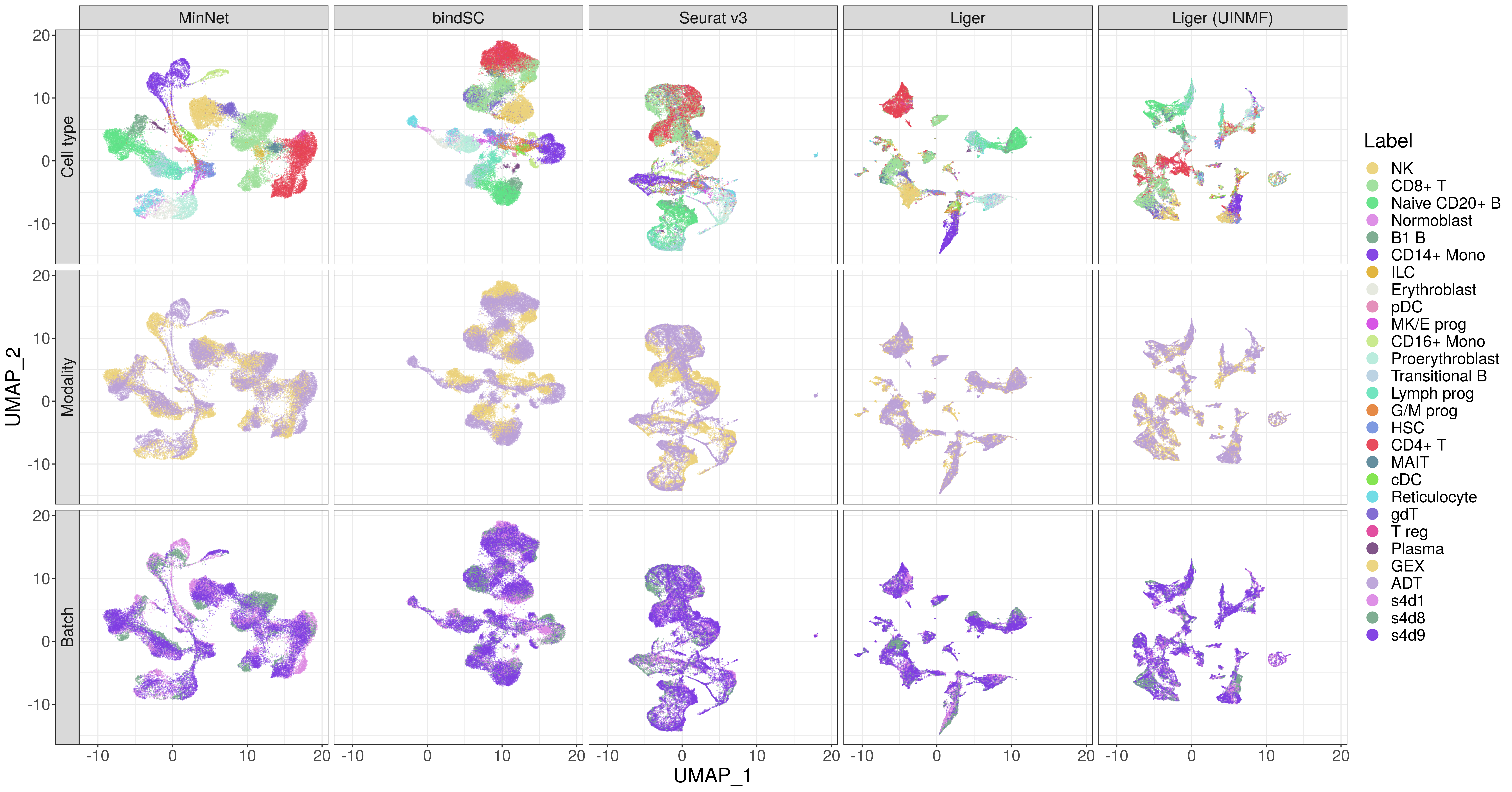

### Supplementary Figure 5

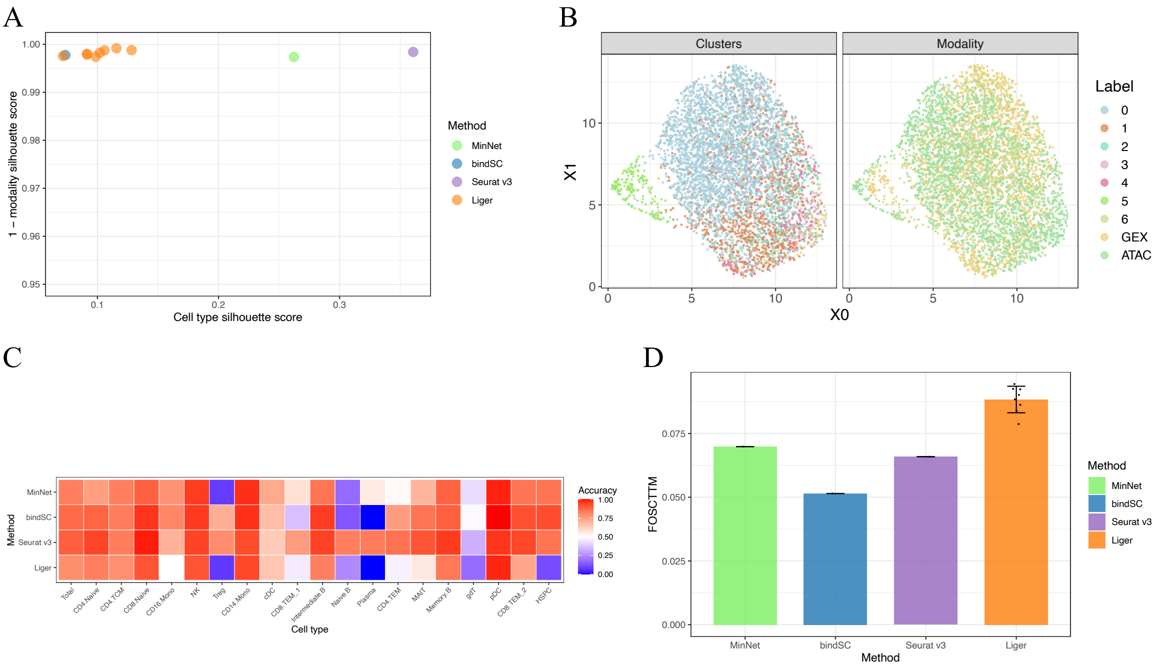

### Supplementary Figure 6

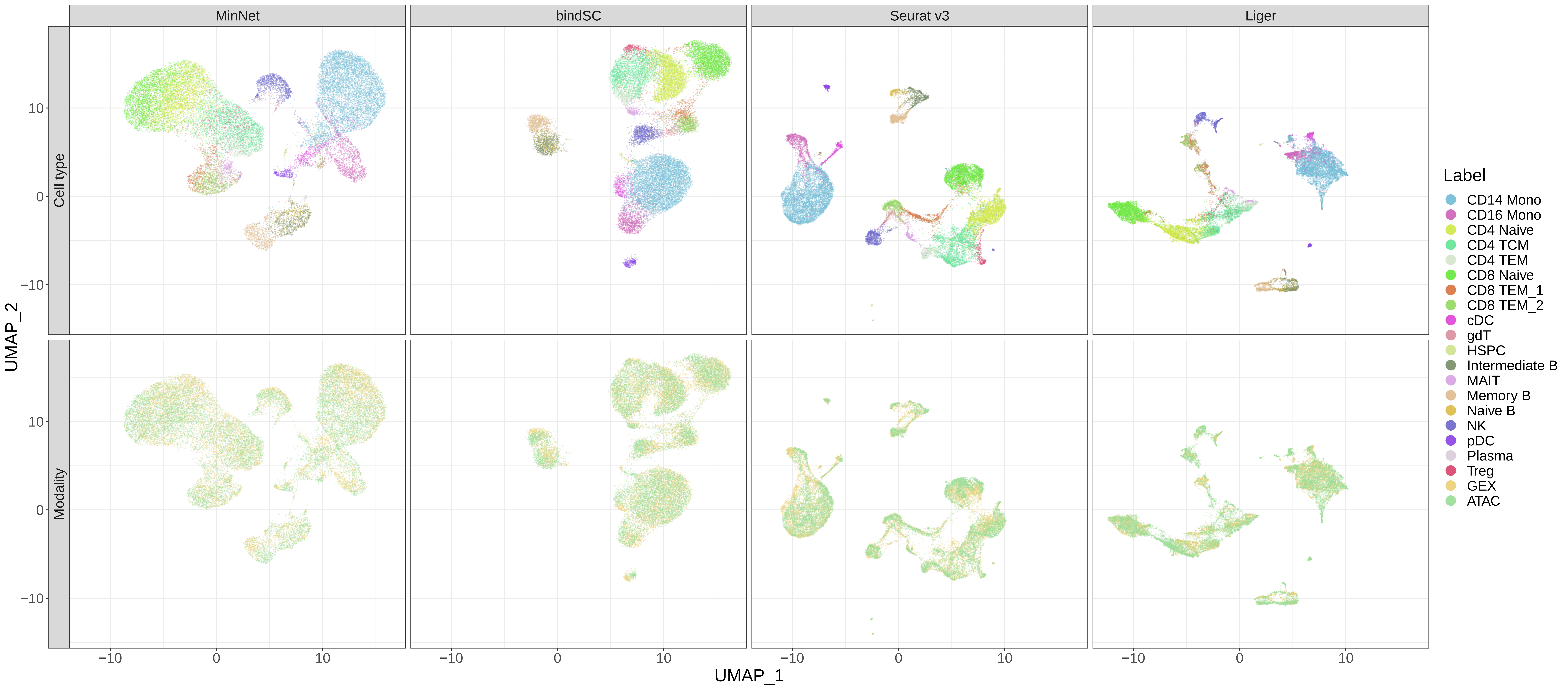

### Supplementary Figure 7

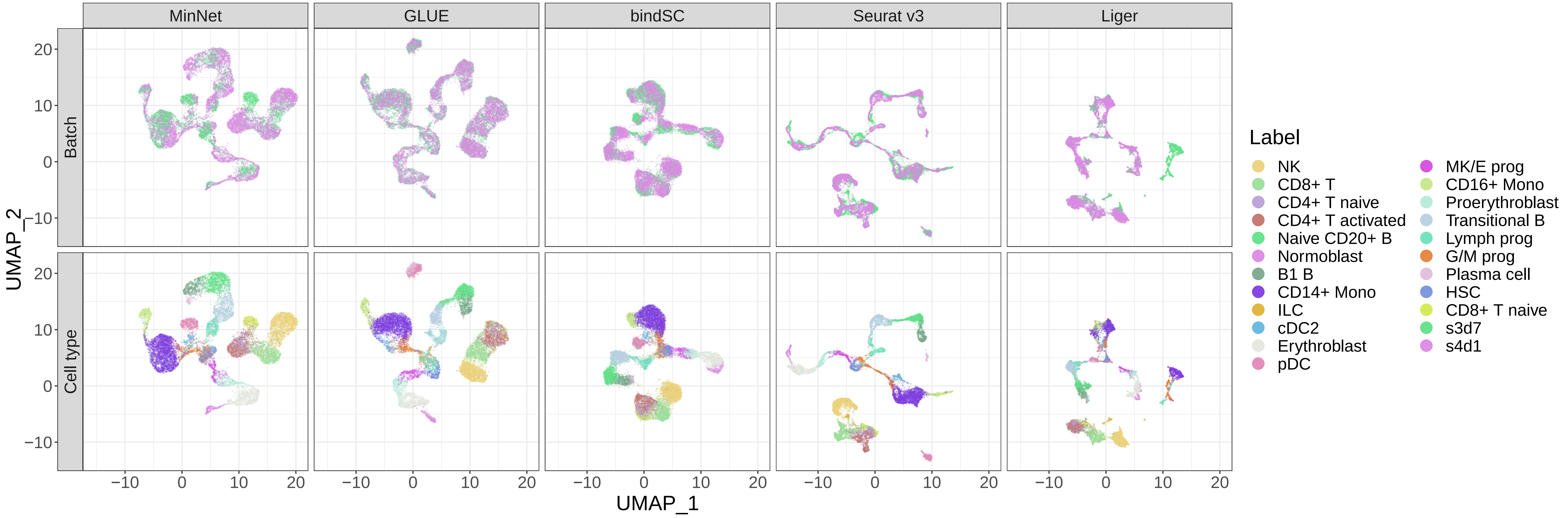

### Supplementary Figure 8

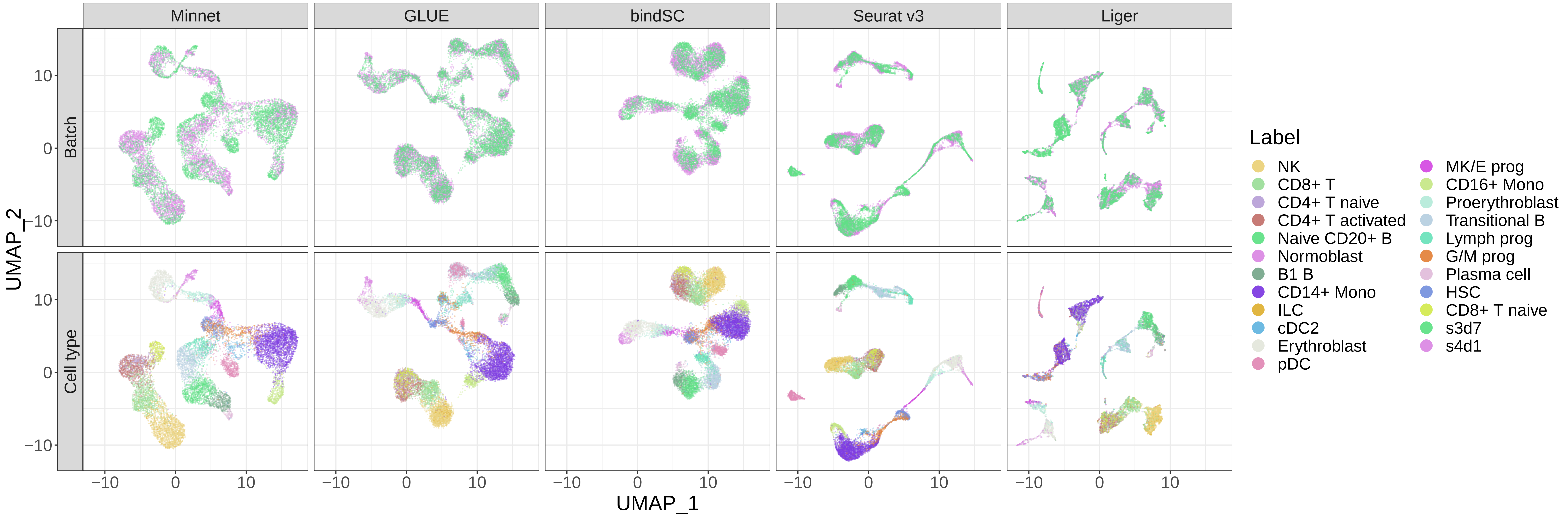

### Supplementary Figure 9

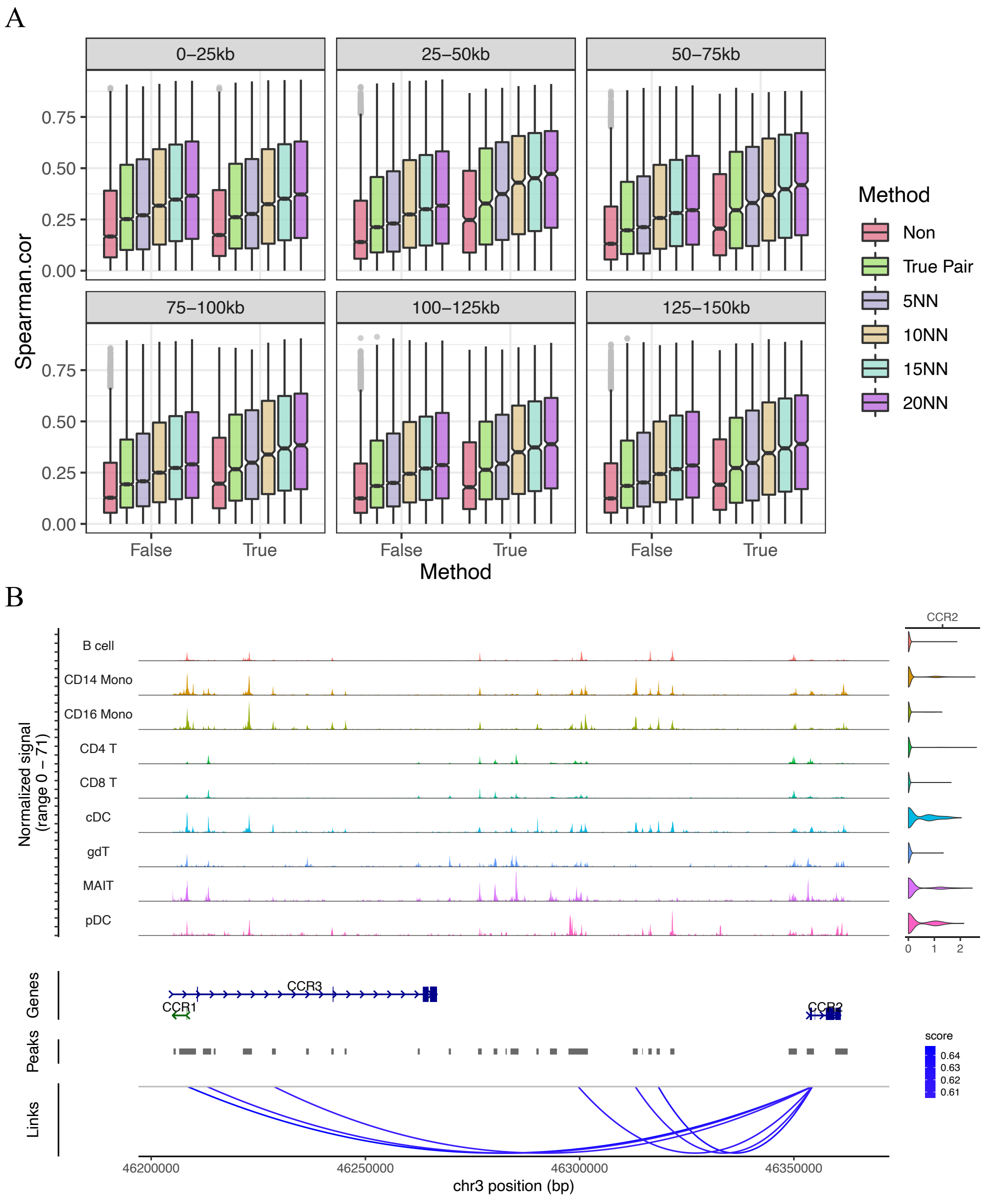

### Supplementary Figure 10

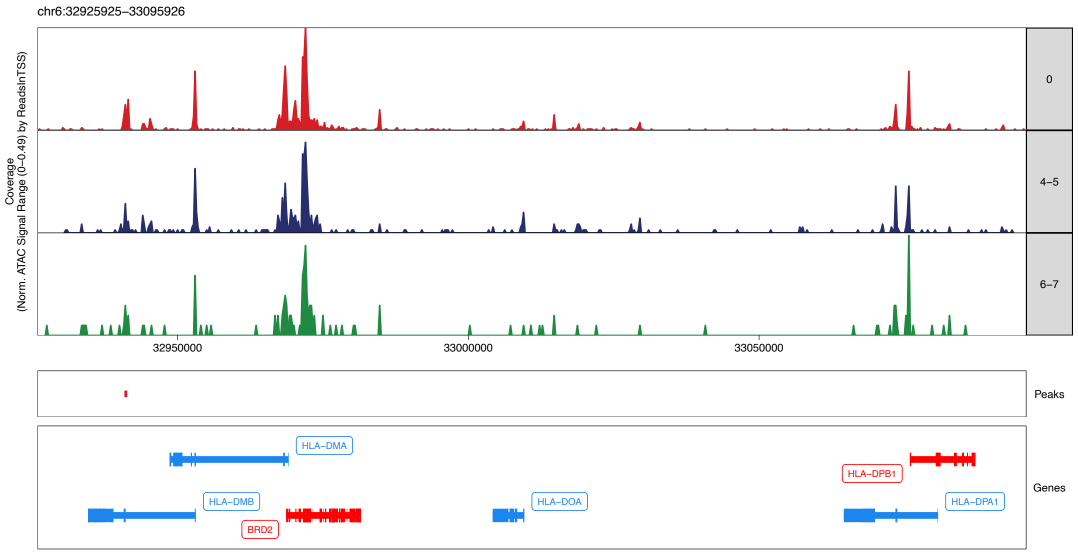

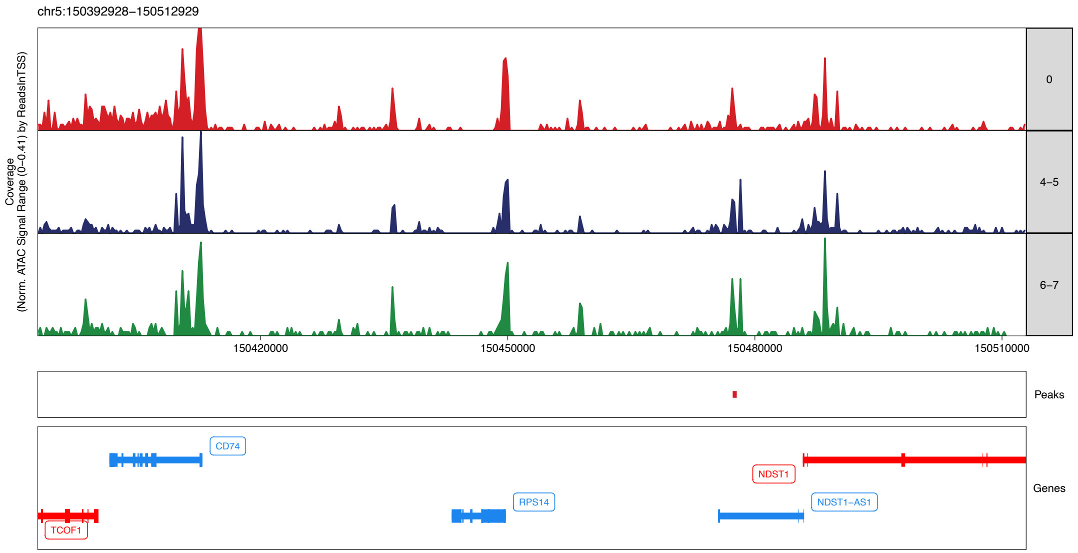
